# Narrowly confined and glomerulus-specific onset latencies of odor-evoked calcium transients in the periglomerular cells of the mouse main olfactory bulb

**DOI:** 10.1101/392274

**Authors:** Ryota Homma, Xiaohua Lv, Tokiharu Sato, Fumiaki Imamura, Shaoqun Zeng, Shin Nagayama

**Affiliations:** Department of Neurobiology and Anatomy, McGovern Medical School at The University of Texas Health Science Center at Houston, Houston, TX, 77030; Britton Chance Center for Biomedical Photonics, Wuhan National Laboratory for Optoelectronics, Huazhong University of Science and Technology, Wuhan, 430074, China; MoE Key Laboratory for Biomedical Photonics, Collaborative Innovation Center for Biomedical Engineering, School of Engineering Sciences, Huazhong University of Science and Technology, Wuhan, 430074, China; Department of Pharmacology, Pennsylvania State University College of Medicine, Hershey, PA, 17033

## Abstract

Odor information is transmitted from olfactory sensory neurons to principal neurons at the glomeruli of the olfactory bulb. The intraglomerular neuronal circuit also includes hundreds of GABAergic interneurons referred to as periglomerular (PG) cells. Stimulus selectivity is well correlated among PG cells that are associated with the same glomerulus, consistent with their highly homogeneous sensory inputs. However, much less is known about the temporal aspects of their activity, including the temporal coordination of their odor-evoked responses. As many PG cells within a glomerular module respond to the same stimulus, the extent to which their activity is temporally aligned will affect the temporal profile of their population inhibitory inputs. Using random-access high-speed two-photon microscopy, we recorded the odor-evoked calcium transients of mouse PG cells and compared the onset latency and rise time among neurons putatively associated with the same and different glomeruli. Whereas the overall onset latencies of odor-evoked transients were distributed across a ~150 ms time window, those from cells putatively associated with the same glomerulus were confined to a much narrower window of several tens of milliseconds. This result suggests that onset latency primarily depends on the associated glomerulus. We also observed glomerular specificity in the rise time. The glomerulus-specific temporal pattern of odor-evoked activity implies that the temporal patterns of inhibitory inputs are unique to individual glomerulus–odor pairs, which may contribute to efficient shaping of the temporal pattern of activity in the principal neurons.

## Introduction

The glomeruli in the olfactory bulb (OB) form an attractive model system for studying signal processing owing to their well-documented neurons and characteristic anatomy, in which the input and output pathways are well segregated. In a glomerulus, sensory inputs from olfactory sensory neurons (OSNs) are transmitted to mitral/tufted cells, which in turn send the signals to higher olfactory centers. Signal processing in the glomerulus involves feedforward excitation and inhibition mediated by hundreds of juxtaglomerular (JG) cells, the diverse interneurons (see below) located in the glomerular layer (GL) (De Saint Jan et al., 2009; Najac et al., 2011; Shao et al., 2012; Carey et al., 2015; Geramita and Urban, 2017). One remarkable trait of these feedforward circuits is that each type of feedforward input (excitatory or inhibitory) to a glomerulus is mediated by numerous JG cells of a specific type. The properties of collective feedforward inputs thus depend on the extent to which the activity of participating neurons is temporally coordinated. However, little is known about the temporal patterns of activity among JG cells that are associated with the same glomerulus.

The term “JG cell” is a generic name for interneurons in the GL and includes multiple types of neurons: major JG cell types include GABAergic periglomerular (PG) cells, glutamatergic external tufted (ET) cells, and GABAergic/dopaminergic superficial short-axon (sSA) cells (Nagayama et al., 2014). PG cells are the most abundant cell type among JG cells. They are monoglomerular (their dendrites project to a single glomerulus) and they receive sensory inputs directly from OSNs or disynaptically from ET cells, depending on the PG cell subtype. ET cells are also monoglomerular and receive direct input from OSNs. These two cell types are the sources of feedforward inhibition and excitation, respectively, in the intraglomerular circuit. The neuronal processes of sSA cells cross multiple glomeruli and are believed to mediate interactions among glomeruli within the GL (Kiyokage et al., 2010). The cellular properties and connectivity of these neurons, including their subtypes, have been extensively studied (Wachowiak and Shipley, 2006; Kosaka and Kosaka 2014, 2016; Burton 2017).

The basic functional properties of JG cells have been revealed previously by *in vivo* recordings (Wellis and Scott, 1990; Tan et al., 2010; Homma et al., 2013; Kikuta et al., 2013; Wachowiak et al., 2013; Fukunaga et al., 2014; Livneh et al., 2014). For example, JG cells fire spontaneously at various rates; this spontaneous activity is phasic and tuned to respiration phase in the majority of cells. In response to odors, JG cell firing increases or decreases and/or shifts in phase. The relationship between glomerular inputs and JG cells has been studied via electrophysiological recordings in genetically tagged glomeruli (Tan et al., 2010), and via calcium imaging with glomerulus-specific labeling (Kikuta et al., 2013). The odor selectivity of JG cells is correlated with that of OSNs as well as with that of other JG cells associated with the same glomerulus. However, neither technique allows simultaneous recording from multiple JG cells at sufficient temporal resolution to analyze how the activity of these cells is temporally coordinated. In this study, we used high-speed two-photon calcium recording (Grewe et al., 2010) to compare the calcium transients in JG cells within and across glomeruli at a high temporal resolution. We demonstrated that the time course of odor-evoked calcium transients is primarily determined by the glomerulus. The onset latency of JG cells was highly heterogeneous, with a ~150 ms difference between the earliest and the latest responses, but onset latency was confined to a much narrower window when we considered only the cells putatively associated with the same glomerulus. Such coordinated activity in JG cells could help to efficiently shape the time course of sensory inputs that are unique to the associated glomerulus.

## Materials and Methods

### Materials

Eight mice (one female) that were the progeny of a Gad2-IRES-Cre mouse (Taniguchi et al., 2011; JAX Stock No. 10802; RRID:IMSR_JAX:010802) and a cre-recombinase-dependent tdTomato reporter mouse (Ai9; Madisen et al., 2010; JAX Stock No. 7909; RRID:IMSR_JAX:007909) were used in this study. An adeno-associated virus (AAV) vector that encodes the GCaMP6f gene under the synapsin promoter (AAV1.Syn.GCaMP6f.WPRE.SV40) was purchased from the UPenn Vector Core. All odorants were purchased from Sigma-Aldrich.

### Animal preparation

All animal procedures were conducted in accordance with an animal protocol that was approved by the Institutional Animal Care and Use Committee (IACUC) of UTHealth.

#### Viral injections

Animals were anesthetized with an intraperitoneal injection of ketamine/xylazine (10/0.5 mg/ml k/x, 12 μl/g bodyweight). The depth of anesthesia was routinely monitored by toe pinches, and additional injections of anesthetic were made to maintain the appropriate depth of anesthesia. Rectal body temperature was maintained between 36.0°C and 37.0°C. The skull above the dorsal OBs was exposed, and two small holes corresponding to two injection sites were made above the posterior end of one OB. For each injection, a glass pipette containing the AAV suspension (no dilution: 9 × 10^12^ GC/ml) was inserted through one of the holes. The pipette approached from the posterior side of the animal, parallel to the A–P axis and tilted 30°–45° from vertical. The tip of the pipette was advanced 250–300 μm from the bulbar surface. A 320–640 nl injection was made at the rate of 64–128 nl/min with a Nanoject II oil-pressure injector (Drummond) in each of the two injection sites. After surgery, mice were left to recover undisturbed for 15–34 days before the recording session. With this injection protocol, uniform labeling of the entire dorsal OB was observed under a fluorescence microscope. In some of the animals (labeled mice 6–8 in the figures), a cranial window was implanted during the surgery for the AAV injections. In these animals, a metal head-plate with a 5 mm round hole (CP-1, Narishige) was attached to the skull with cyanoacrylate glue and dental acrylic. A round cranial window (~3 mm in diameter) that covered both dorsal bulbs was made through the hole of the head-plate. The dura mater was not removed. The exposed bulb was sealed with 0.8% agarose and a 3 mm round coverslip (CS-3R, Warner Instruments), which was cemented with cyanoacrylate glue and dental acrylic.

#### Data acquisition

Animals previously injected with AAV were anesthetized with an intraperitoneal injection of urethane (6% w/v, 20 μl/g bodyweight for induction, with the depth of anesthesia monitored by toe pinches and maintained by additional injections). Animals breathed freely throughout the experiment, and rectal body temperature was maintained between 36.0°C and 37.0°C. Animals with a pre-installed cranial window (see above) were transferred to the microscope stage once the level of anesthesia was stable. For animals without a pre-installed cranial window, a metal head-plate was attached to the skull with cyanoacrylate glue and dental acrylic. Then, a craniotomy was made in the AAV-injected side of the OB and the opening was filled with 1.2% agarose and sealed with a pre-cut piece of coverslip, which was cemented with cyanoacrylate glue. Following the cranial window preparation, the animal was transferred to the microscope stage. The microscope stage was equipped with an angle-adjustable stage (MAG-3, Narishige) to allow fine adjustments of the angle of the bulbar surface relative to the objective. Breathing was monitored with a piezo-electric sensor placed beneath the chest, and the signal was recorded together with the optical signals.

### Optical recordings

All optical recordings (i.e., wide-field and two-photon) were conducted with the same upright fluorescence microscope (Olympus) combined with a motorized stage (Scientifica). All optics were implemented on a vibration-isolating air table (Newport). For wide-field imaging, the excitation light was provided by a 470 nm LED module (M470L2, Thorlabs). The filter set was a standard GFP cube (GFP-4050A-OMF-ZERO, Semrock; 466/495/525 nm exciter/DM/emitter). A 10×/0.3 NA objective lens (Olympus) and a 0.35× tube lens (Olympus) were used, and the images were captured by a high-speed CCD camera (NeuroCCD-SM256, RedShirtImaging) at 125 Hz for 12 s. With this configuration, a 1.75 × 1.75 mm^2^ area of bulbar surface was imaged at 128 × 128 pixels. For two-photon recordings, a custom-built acousto-optic deflector (AOD)-based two-photon scanner that is capable of random-access recording was used (Iyer et al., 2006; Grewe et al., 2010). Random-access scanning is a mode of two-photon recording that allows one to record from only an arbitrary set of pre-designated pixels in the field of view, typically at a much higher sampling rate than conventional full-field imaging. A random-access pattern scanning mode was adopted (Grewe et al., 2010), in which a fixed set of adjacent pixels were also sampled in every sampling cycle so that the total light exposure time for each pixel was reduced. The detailed design of the AOD two-photon scanner is described elsewhere (Lv et al., 2006). The light source was a Ti:Sapphire laser (MaiTai HP DS, Spectra-Physics), and 920 nm light was used for all two-photon recordings. The emitted fluorescence was routed via a dichroic mirror (FF665-Di02, Semrock) to the detector module, which was located adjacent to the filter cube to minimize the length of the light path. The detector module contained an IR-block filter (FF01-680/SP, Semrock) as well as a 562 nm dichroic mirror (FF562-Di02, Semrock) that separates shorter (green) and longer (red) wavelength light. The separated shorter and longer wavelengths of light were detected by a GaAsP photosensor module (H7422A-40, Hamamatsu) and a photomultiplier tube (R1924A, Hamamatsu), respectively. Full-frame scan recordings for functional imaging were made at 2.1 Hz for 15 s with a resolution of 217 × 217 pixels. With the Olympus 25×/1.05 NA objective lens that used in this study, the field of view was 219 × 219 μm^2^. For the high-speed random-access recordings, 15 sites were pre-determined on the basis of the odor-response data obtained during the frame scan recordings (see Fig. 2B). Although the system is capable of recording a larger number of sites (at a lower sampling rate), 15 was about the maximum number of cells with clear odor-evoked responses that could be identified from a given field of view. Therefore, all random-access recordings presented in this study were conducted with 15 sites, each of which covered a single neuron, at a sampling rate of 667 Hz for 15 s.

**Figure 1.**
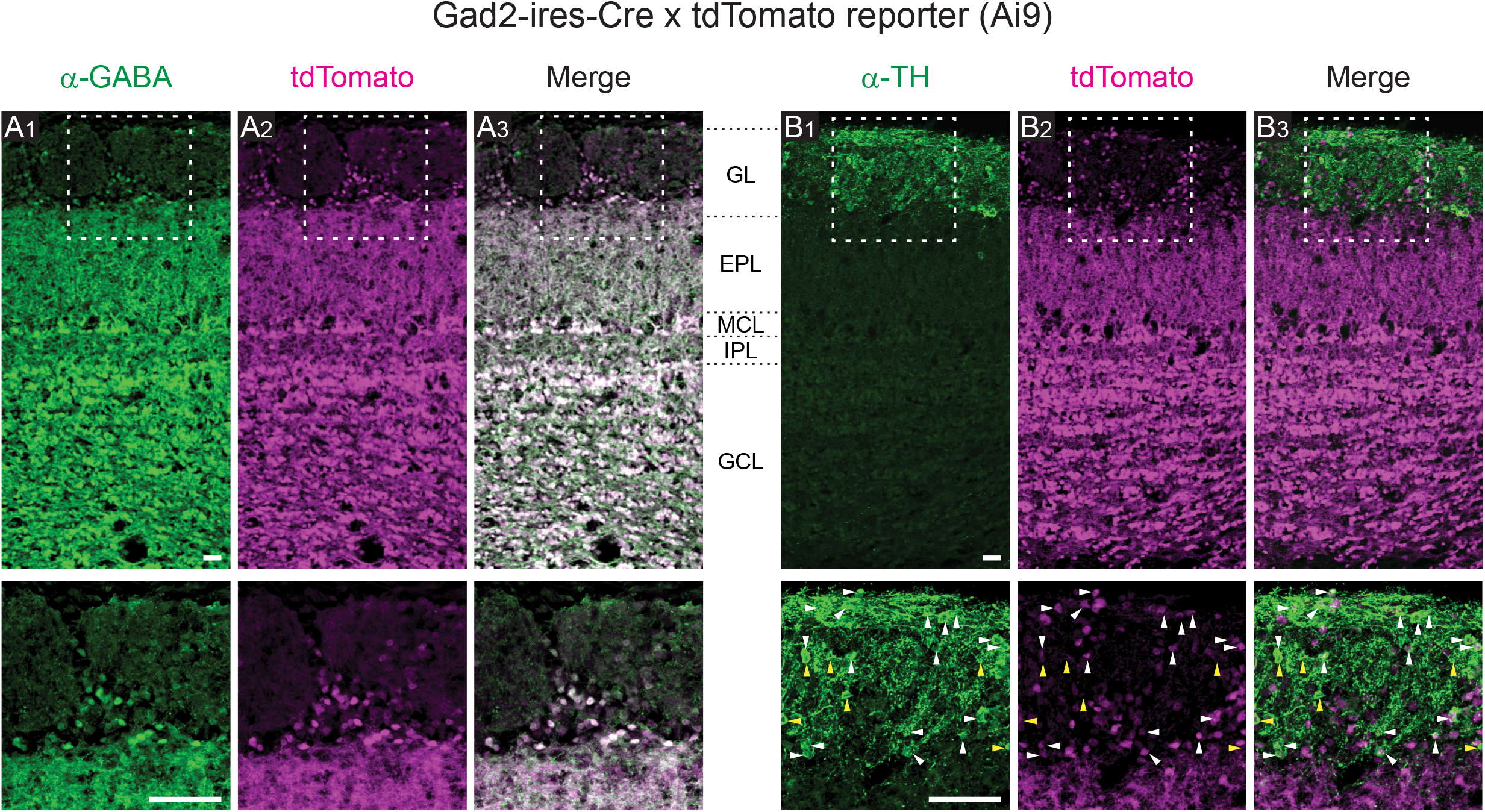
Expression patterns of tdTomato. The progeny of Gad2-IRES-Cre and cre-dependent tdTomato reporter mice were used in this study. A1–3. Spatial pattern of immunolabeled GABA (A1), tdTomato (A2), and the merged image (A3). The region indicated by a white box in the top panels is presented at a higher magnification in the corresponding bottom panels. B1–3. Spatial pattern of immunolabeled TH (B1), tdTomato (B2), and the merged image (B3). White and yellow arrowheads indicate examples of TH^+^/tdTomato^+^ cells and TH^+^/tdTomato^−^ cells, respectively. Approximate positions of layer boundaries are indicated by black dotted lines in the space between the panels A3 and B1. Scale bars: 50 μm.

**Figure 2.**
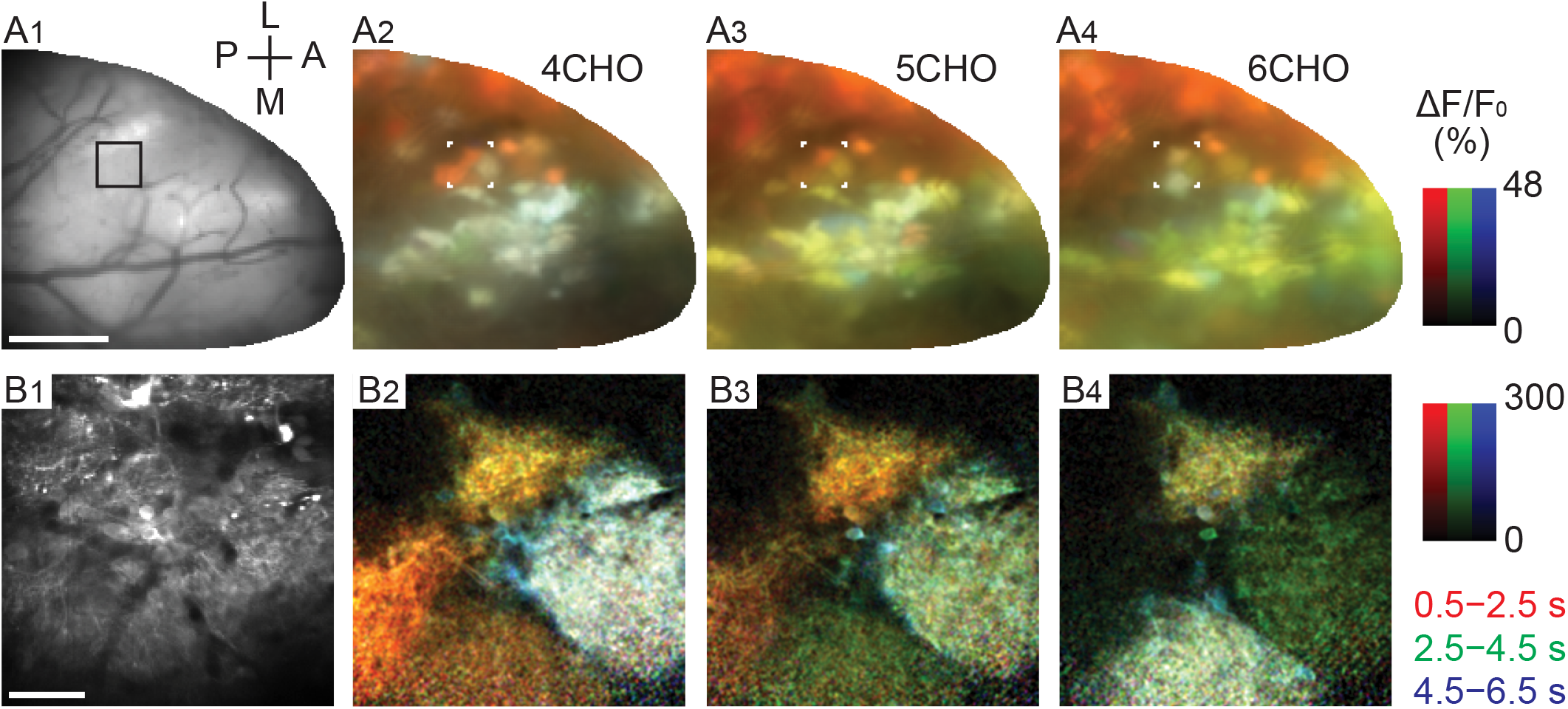
Activity (ΔF/F_0_) maps color-coded by the phase of response in wide-field and two-photon imaging. A1. Surface image for wide-field (single-photon) imaging. Scale bar: 500 μm. A2–4. Maps of odor-evoked response to 4CHO, 5CHO, and 6CHO, respectively. Each map is a synthesis of red, green, and blue maps that represent the periods 0.5–2.5, 2.5–4.5, and 4.5–6.5 s after stimulus onset, respectively. B1. A two-photon image of the selected area, indicated by the black box in A1 and by white corners in A2–4. Scale bar: 50 μm. B2–4. Maps of odor-evoked response, with the colors representing the same periods as in A2–4.

### Stimulus presentation

Odor stimuli were presented via a custom-built olfactometer, which was described previously (Kikuta et al., 2013). Briefly, a mixture of odorous gas (diluted odorant bubbled with 100% nitrogen) at 100 ml/min and carrier gas (filtered air) at 400 ml/min was continuously supplied but continuously vacuumed near the tip of the applicator. The vacuum was suspended with an electric command signal, resulting in the presentation of odorous gas until the vacuum resumed. In a test conducted with a photo-ionic odorant sensor (mini PID model 200B, Aurora Scientific), the onset and offset of actual odorant delivery was delayed by approximately 100 ms relative to the timing of the command signals. The olfactometer was controlled by a custom-written program in LabVIEW (National Instruments; RRID:SCR_014325) that enabled the stimulus presentation to be synchronized to the respiration cycle of the mouse. Four seconds after the onset of data acquisition, the command signal for an odor stimulus was sent at the peak of the first respiration signal (corresponding to the end of an inhalation cycle) so that the odorant did not reach the mouse in the middle of an inhalation. The duration of the stimulus was either 2 s or a single respiration cycle, in which case the command signal to terminate the odor stimulus was sent at the peak of the respiration signal in the immediately subsequent cycle.

For the odor stimuli, five single-molecule odorants that were aliphatic aldehydes with carbon chain lengths of three to seven (referred to as 3–7CHO, respectively) were used. All odorants were diluted 1:100 with mineral oil and further flow-diluted 1:5 at the olfactometer. Therefore, the nominal final concentration of the odorants was 0.2%. Five odorants (3–7CHO) were presented in a block design, with each odorant presented once in a block of five trials. For mice 1–5, the odorant presentation order was the same in every block. For mice 6–8, the order was pseudo-randomized with the constraint that the same odorant was not presented in two sequential trials. In alternate blocks, the stimulus was presented for 2 s or for a single respiration cycle.

### Immunohistochemistry

Mice were deeply anesthetized and fixed by transcardial perfusion with 4% paraformaldehyde in 0.1 M sodium phosphate buffer (pH 7.4). Then, brain samples were collected and post-fixed overnight at 4°C. The samples were cryoprotected in 30% sucrose (w/v) in PBS (pH 7.4) and embedded in optimal cutting temperature compound (Sakura Finteck). The olfactory tissues were cut on a cryostat into 20 μm sections (coronal) and stored at −20°C until use. The slices were pretreated in TBS-T (10 mM Tris-HCl (pH 7.4), 100 mM NaCl with 0.3% Triton-X100 (v/v)) for 15 min and blocked with blocking buffer (5% normal donkey serum (v/v) in TBS-T) at 20–25°C for 1 h. The slices were then incubated with primary antibodies diluted in blocking buffer overnight at 4°C. Sections were washed with TBS-T, then incubated with secondary antibodies for 1 h. The immunoreacted sections were washed and mounted with FLUORO-GEL mounting medium (Electron Microscopy Sciences, #17985-11).

The following antibodies were used. Primary antibodies: anti-GABA (MilliporeSigma, #A2052, rabbit, 1:100) and anti-TH (MilliporeSigma, #AB152, rabbit, 1:500). Secondary antibody: Alexa Fluor 488 donkey anti-rabbit IgG (Thermo Fisher Scientific, #A21206, 1:300).

Six to eight serial images were captured at intervals of 1.5 μm with an Olympus FluoView FV1000 laser scanning confocal microscope with a 20× objective lens (UPLSAPO 20×), with or without 4× digital zoom. Then, the acquired images were converted to a Z-stack with the software FV10-ASW viewer (Olympus).

### Data analysis

There were minor differences in the experimental conditions (e.g., the timing of the craniotomy and the order of stimulus presentation; see above) between the two subsets of mice (mice 1–5 versus mice 6–8). Since no difference was noticed between the data from these two subsets, they were not discriminated in any of the following analyses. All data analyses and preparation of figures were carried out in Fiji/ImageJ (Schindelin et al., 2012; RRID:SCR_002285) and MATLAB (MathWorks, RRID:SCR_001622) with custom scripts. In the recordings in this study, no prominent photobleaching of fluorescence was observed within a trial, and thus no correction for photobleaching was made unless explicitly described.

#### Spatiotemporal activity maps

For wide-field imaging, spatial patterns of odor-evoked activity were visualized as the difference between images from the pre-stimulus period and the response period (i.e., ΔF/F_0_ maps; Fig. 2A). A baseline image from the pre-stimulus period was obtained by averaging the frames from 3 to 4 s after acquisition onset. Images for three different response periods (early, middle, and late) were obtained by averaging the frames in the 2 s time windows following 4.5, 6.5, or 8.5 s, respectively, from the acquisition onset (0.5, 2.5, 4.5 s from the stimulus onset). In this specific analysis, the trial-by-trial jitter of inhalation onset or stimulus onset was not taken into account. Then, the pre-stimulus-period image was subtracted from each of three response-period images. The subtracted images were divided by a spatially filtered pre-stimulus-period image (two passes of a 3 × 3 mean filter) for normalization. After the division, the resulting image was also filtered by two passes of a 3 × 3 mean filter. Red, green, and blue color channels were assigned to the resulting images for the early, middle, and late response periods, respectively, and then all three channels were merged to form a full-color map. Therefore, the brightness of the resulting activity maps reflects the response amplitude and the hue reflects the character of the response time course. The intensity scale was set to the same value in each of the three color channels. Essentially the same procedure was applied to obtain the activity maps for two-photon full-frame imaging (Fig. 2B).

#### Basic signal processing of the time-course data

Data from two-photon random-access recordings are not a series of images, but a set of time series that resemble the data from multi-electrode recordings. Thus, no image processing is involved in the analysis. Time courses were converted to ΔF/F_0_ by subtracting the averaged data from the first 1% of data points (150 ms; F_0_) and then dividing by F_0_. A box filter with a 100 ms window was applied to all the time courses unless specified.

#### Response-based grouping of cells

The subset of recorded cells that surrounded the same glomerulus had highly correlated odor selectivity (Figure 3). Under the assumption that these neurons were associated with the same glomerulus, such neurons were grouped using the following two criteria. First, the neurons were directly adjacent to the glomerulus of interest and no other glomerulus was located in between. Second, the response profile of a given neuron was closer to that of all the in-group recorded neurons than to that of any out-group recorded neurons. In other words, the least similar in-group neuron had a more-similar response profile than the most-similar out-group neuron. Cosine similarity (cosine of the angle between two vectors) was used as a measure of the similarity between two response profiles. A response profile here is defined as a vector representing the response amplitudes to the five different odors, or a vector representing all pairwise differences (ten pairs from five odorants) of response amplitudes. When at least one of these two response profiles satisfied the above criteria, the neuron was considered as qualified. Response amplitudes were defined as the integral of the ΔF/F_0_ trace (area under the trace) from the trials with single-respiration-cycle stimulation. In this analysis, since even tiny responses are potentially influential, photobleaching/drift was corrected for by subtracting a straight line that was best fitted to the 0–4 s (pre-stimulus period) and the 14–14.5 s period (post-response period) of the time course, only when the average of the latter period had a negative value. The correction was made only in the trials with downward changes because (1) prominent photobleaching/drift was observed only in the negative direction, and (2) applying correction to positive values would significantly distort the subset of data that showed long-lasting odor responses.

**Figure 3.**
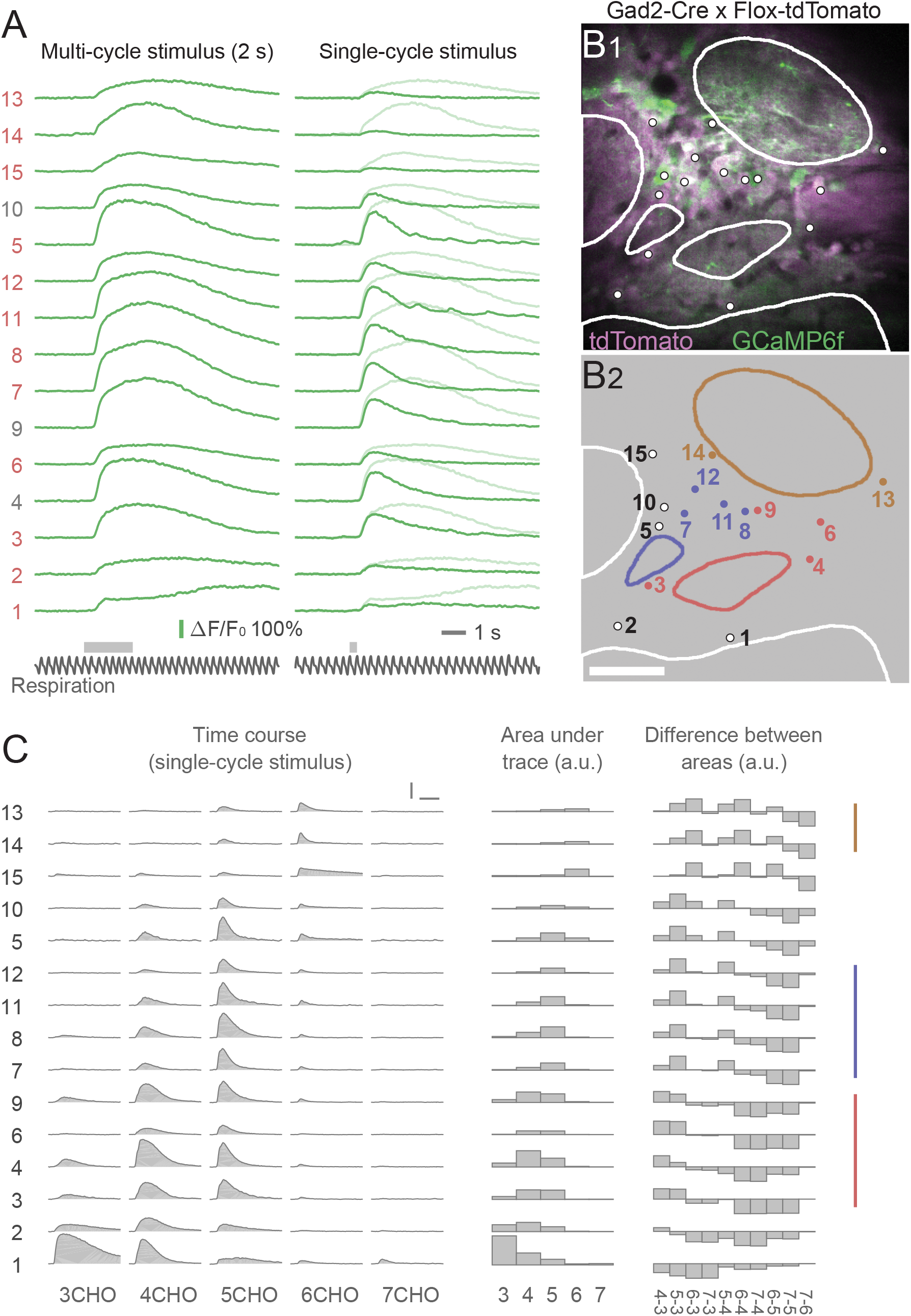
Odor-evoked calcium transients and grouping of JG cells by response profile. A. An example data set from a single trial of random-access scanning of 15 ROIs (cells). Left traces are responses to a 2 s odor stimulus, and right traces are responses to a single-respiration-cycle stimulus. The bottom black traces show the respiration signal. Gray horizontal bars above the respiration signals indicate the timing of valve opening for odor presentation. Note that actual odor presentation lags approximately 0.1 s behind the valve opening. The ordering of the ROIs is intentional, based on the result of grouping shown in C. B. Two-photon image of the recording site (B1) and the ROI indexes (B2). Dots and contours represent the ROIs and glomeruli, respectively. Non-white colors of dots and contours in B2 indicate the groups presented in C. Scale bar: 50 μm. C. Odor-evoked responses of the same ROIs to five odors (single-cycle stimulus). The left block shows the response time courses. The middle block shows the areas under the time courses as bar charts. The right block shows the difference in areas between every possible pair of odors. Colored vertical lines at the right indicate the groups of cells putatively associated with the same glomerulus (see text for details). Scale bars: 3 s (horizontal), 100% ΔF/F_0_ (vertical).

#### Determining the onset latency and rise time of calcium transients

The onset latency of an odor-evoked calcium transient was defined via the following procedure (see also a visual explanation in Fig. 5A). All ΔF/F_0_ traces were preprocessed with a 50 ms box filter. For a given trace, the baseline period was defined as a 100 ms time window immediately before the stimulus onset (the onset of the command signal). The baseline ΔF/F_0_ level was determined as the mean value over the baseline period. To determine the noise level, the 4 s pre-stimulus period was divided into eight 0.5 s blocks, calculated the standard deviation within each block, and then selected the minimum among these. This procedure effectively eliminated the contribution of the occasional calcium transient (see Fig. 4A) on the noise level. The provisional onset latency was determined as the first time point in which 95% of the data points in the subsequent 100 ms exceeded the threshold (2.5 times the noise level). Then, the trace within the subsequent 100 ms time window was fitted to a straight line. The intersection of the fitted line and the baseline level was considered the onset of the calcium transient. The onset latency was defined as the duration from the onset of the first inhalation with the odor stimulus to the onset of the calcium transient. Although this procedure determined the onset time more accurately than simple threshold methods in high signal-to-noise (s/n) ratio data, it frequently resulted in obviously wrong values when the s/n ratio was lower.

**Figure 4.**
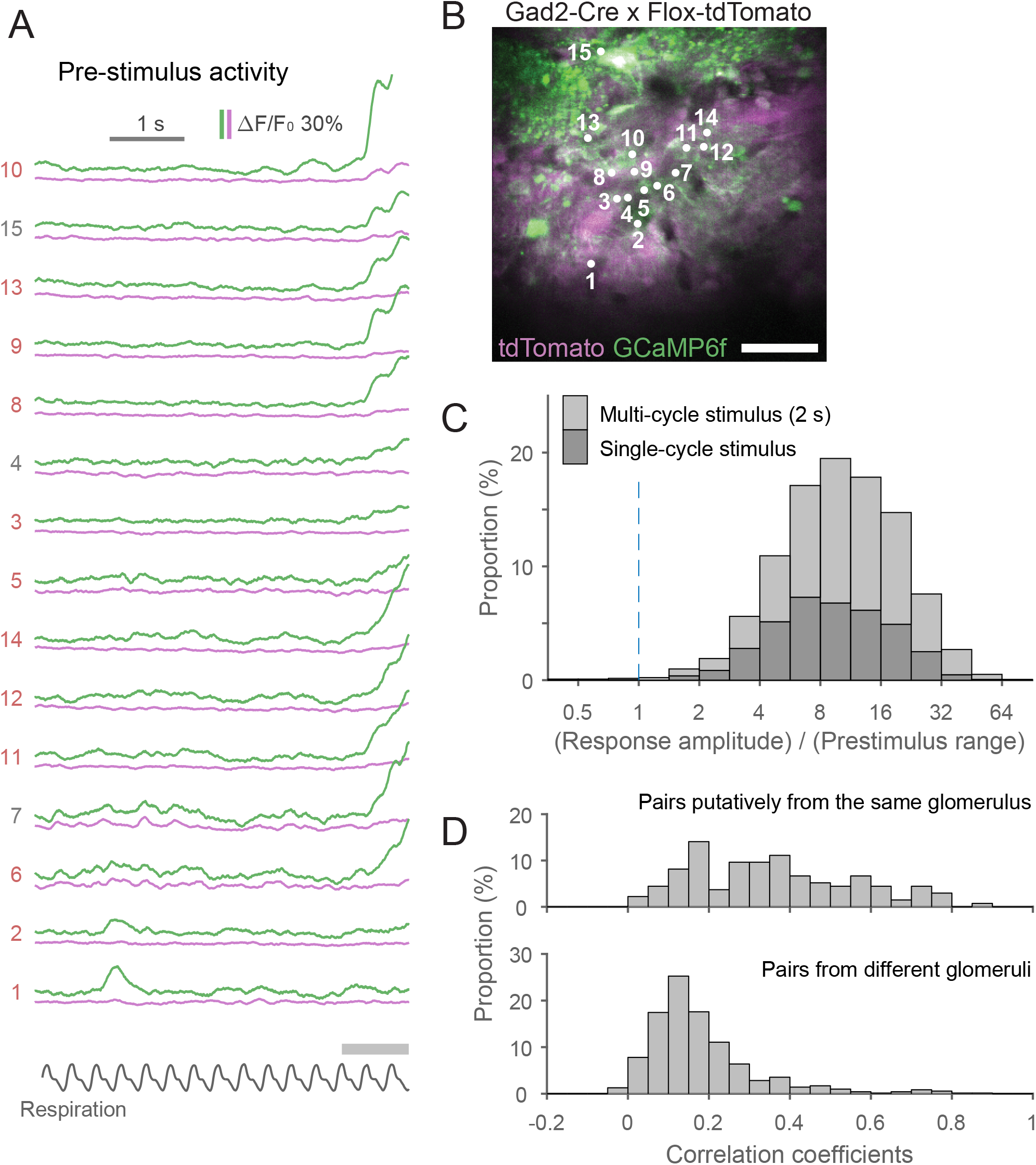
Characterization of calcium signals in the pre-stimulus period. A. An example data set from the pre-stimulus period in a trial. Green traces are the signal from the green channel (GCaMP6f), and magenta traces are the signal from the red channel (tdTomato). Apparent odor-evoked changes in some of the magenta traces (e.g., ROI 10) suggest a minor contribution of the GCaMP signal to the red channel. The relative contribution from the GCaMP signal may vary depending on the relative expression levels of GCaMP and tdTomato at an individual ROI. B. Two-photon image of the recording site with ROIs and their indexes. Scale bar: 50 μm. C. Stacked histogram of the ratio between peak response amplitude and range (the distance between the 1st and 99th percentiles) in the pre-stimulus period. Note the logarithmic scale on the x-axis. The response is much larger than the fluctuations in the pre-stimulus period in the vast majority of cases. D. Histograms showing the distribution of pairwise correlation coefficients for the time courses of activity during the pre-stimulus period. Upper and lower histograms show data from cell pairs putatively associated with the same glomerulus and pairs from different glomeruli, respectively.

**Figure 5.**
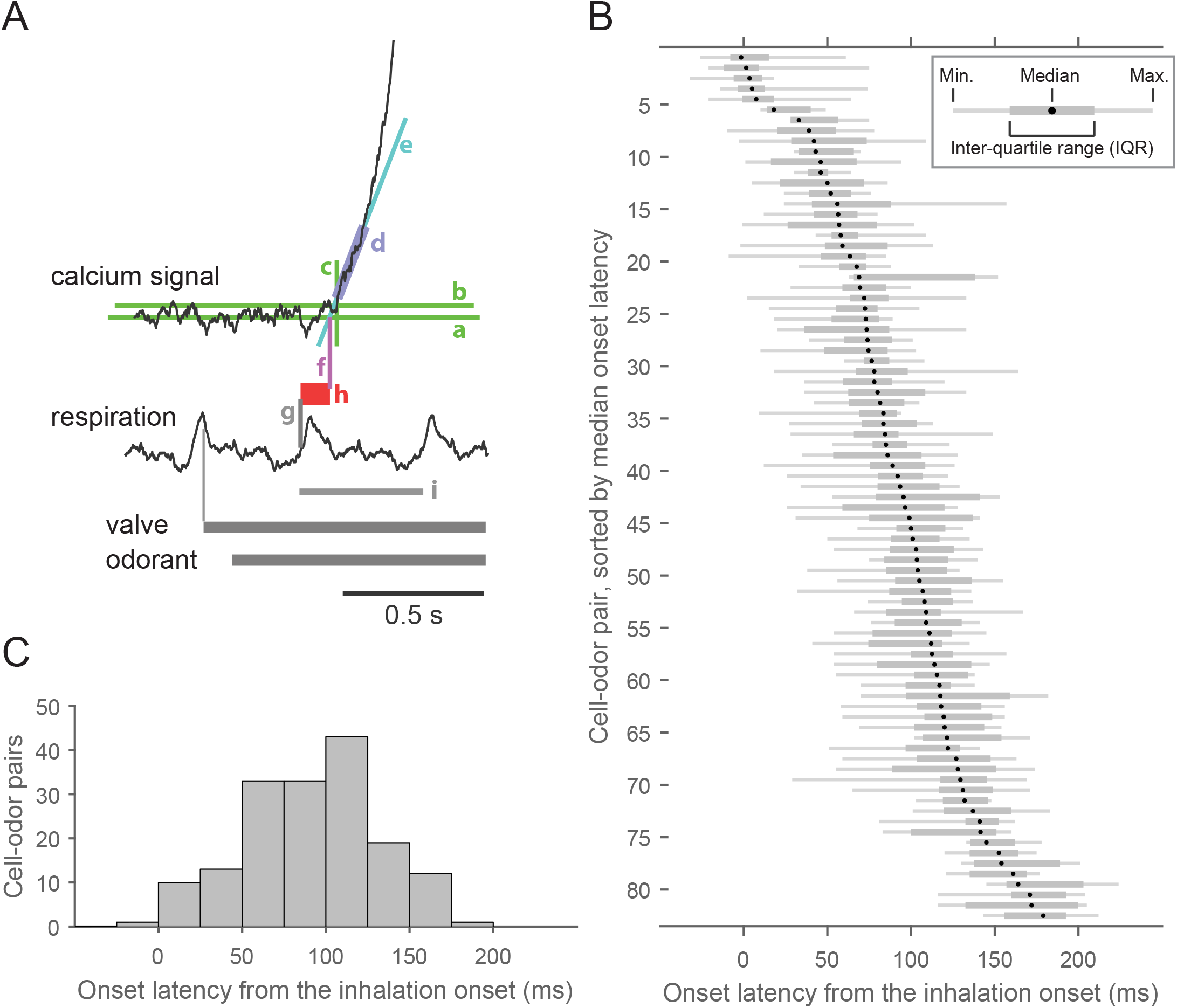
Onset latencies of odor-evoked calcium transients are heterogeneous across JG cells. A. Graphic representation of the definition of onset latency. (a) Baseline, defined as the mean of the pre-stimulus period signal. (b) Threshold, defined as 2.5 times the standard deviation of the pre-stimulus period signal. (c) Time point at which the signal exceeds the threshold. (d) Time window of the first 100 ms above the threshold. (e) Regression line of the signal in the 100 ms time window. (f) Point where the regression line crosses the baseline. This time point is considered the onset of the calcium transient. (g) Onset of the first inhalation with the odor stimulus. (h) Onset latency, defined as the distance between the onset of inhalation and the onset of the calcium transient. B. Distribution of onset latencies. Each row represents a single cell–odor pair as a box-and-whisker plot (see inset). cell–odor pairs are arranged according to median onset latency for clarity (83 cell–odor pairs from 8 recording sites, out of all 165 pairs, are presented). Each box-and-whisker plot represents data from 5–17 trials. C. Histogram showing the distribution of medians shown in B.

Therefore, for this analysis, only data with a relatively high s/n ratio were considered. The s/n ratio was determined as the slope of the fitted line (in ΔF/F_0_ per s) divided by the noise level defined above (in ΔF/F_0_). The threshold s/n ratio was determined empirically and set to 40 (s^−1^). In addition, in a small fraction of trials, the onset of odor-evoked calcium transients accidentally coincided with a “spontaneous” calcium transient. Such trials were discarded. The estimated onset latency was used for further analysis (presented in Figs. 5–7) when the latency was successfully estimated in three or more trials (out of 5–17 trials) for a given cell–odor pair.

**Figure 6.**
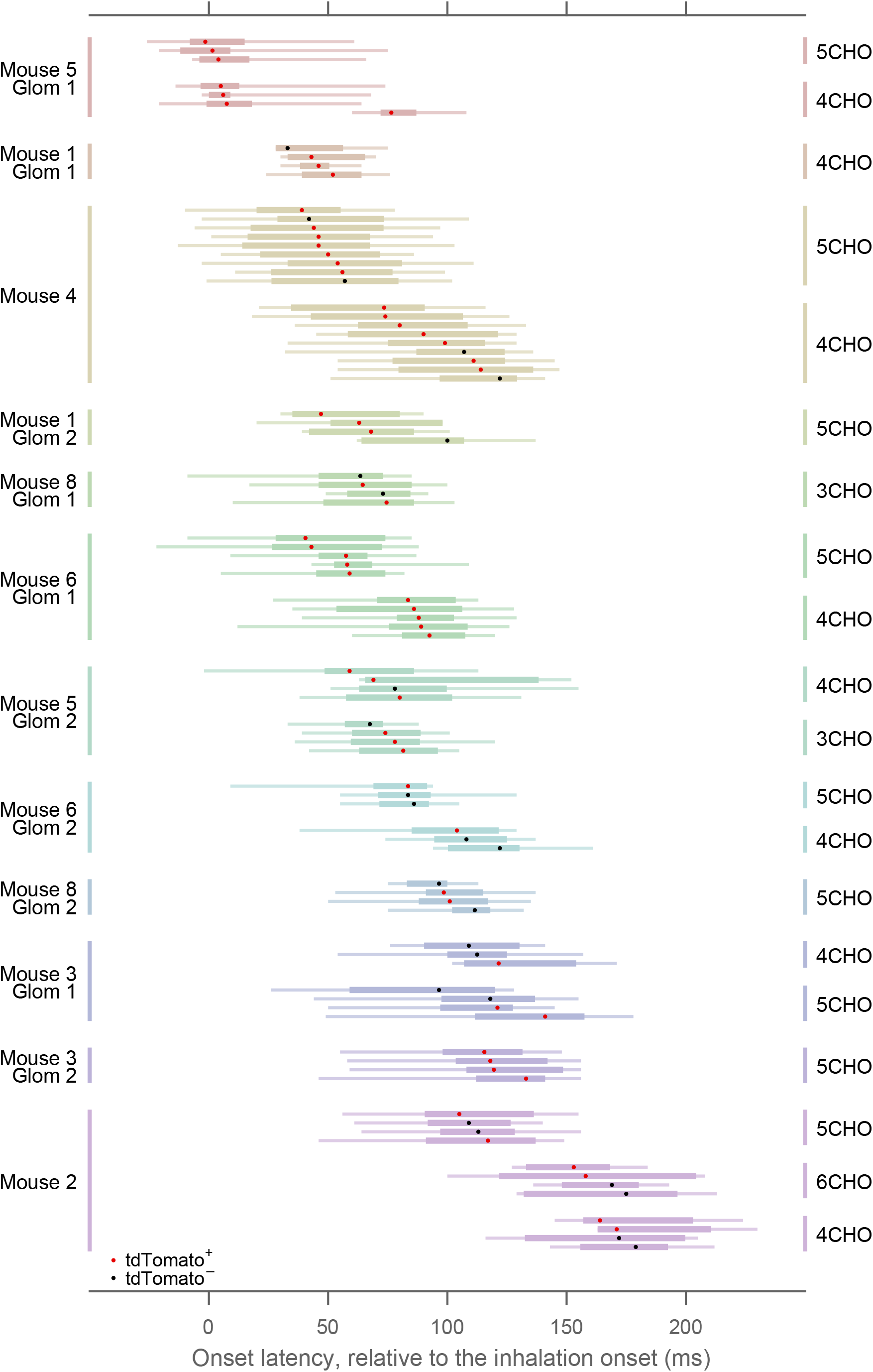
JG-cell onset latency strongly depends on the putative glomerular association. A subset of the box-and-whisker plot from Fig. 5 is presented, rearranged according to glomerulus–odor pairs. Different glomeruli are presented in different colors. In some of the glomeruli, data from more than one odorant were available. Dots in the box-and-whisker plots represent the median, and expression of the GAD2 marker tdTomato is indicated by the dot color (red for tdTomato^+^ and black for tdTomato^−^). Mouse and glomerulus identity are at the left. Odorant is indicated at the right.

**Figure 7.**
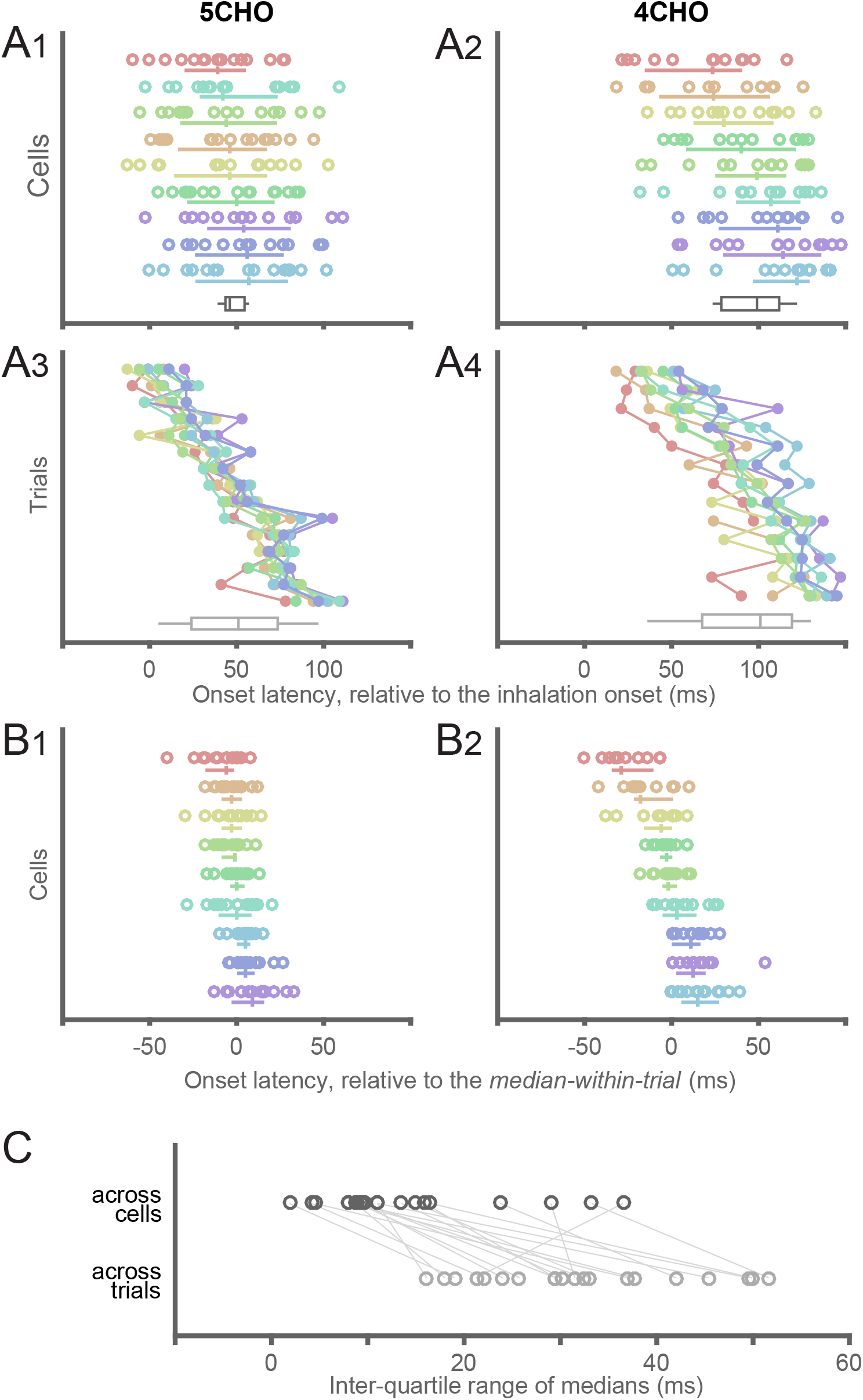
Detailed analysis of onset latency across cells putatively associated with the same glomerulus. The onset latencies of cell–odor pairs from a glomerulus in mouse 4 (see Fig. 6) are presented with an alternative visualization. A1–2. Reconstruction of plots in Fig. 6, except that each trial was explicitly plotted. Each row represents an individual cell, as in Fig. 6. The median and inter-quartile range for each cell are presented as the accompanying vertical and horizontal lines, respectively. Colors represent individual cells and are preserved across all graphs in panels A and B. The cells are sorted by their median onset latency. Left and right graphs show the responses to two different odors. The black box-and-whisker plots at the bottom of each panel show the distribution of median onset latency across cells. A3–4. The same data as in A1–2, but rearranged so that each row represents an individual trial. Note that the variances in individual rows are markedly reduced. Trials are sorted according to their median onset latency, not by the order of acquisition. The gray box-and-whisker plots at the bottom of each panel show the distribution of median onset latency across trials. B. Onset latencies aligned to the median onset latency within the trial for each data point. C. Inter-quartile ranges (IQRs) of median onset latency across cells (black box-and-whisker plots in A1–2) and across trials (gray box-and-whisker plots in A3–4) are compared for all glomerulus–odor pairs. IQRs across cells are smaller in nearly all cases.

The rise time of an odor-evoked transient was defined as the duration between the time points when the signal reached 20% and 80% of the peak amplitude. For this analysis, all ΔF/F_0_ traces were preprocessed with a 100 ms box filter. The baseline level was defined in the same way as the protocol for determining onset latency (see above). The peak amplitude was determined as the maximum value of the moving average of a 100 ms time window. Time points at 20% and 80% of the peak amplitude were defined as the earliest time point when half of the data points exceeded the threshold value in a 100 ms time window centered at the time point being considered.

To compare the onset latency and rise time between tdTomato^+^ and tdTomato^−^ neurons, taking into account their glomerular association, relative onset latency and relative rise time were used. Relative latency, for example, was defined for each group of cells putatively associated with the same glomerulus (including both tdTomato^+^ and tdTomato^−^ neurons). The relative latency ranged from zero (shortest) to one (longest), with equal spacing among neurons in the same group. For instance, if a group contained five neurons, their relative latencies would be 0, 0.25, 0.5, 0.75, and 1, from shortest to longest onset latency. If tdTomato^+^ neurons have shorter or longer onset latencies on average, the distribution of relative onset latencies would be different in tdTomato^+^ and tdTomato^−^ neurons. The same method was applied to rise times.

### Experiment design and statistical analyses

Eight Gad2-tdTomato mice were used (seven males and one female (mouse 2); 12–32 weeks old at the time of recording). Imaging experiments were carried out in one bulb from each animal. The glomerulus-level analysis of onset latency (Fig. 6) includes data from 12 glomeruli in 7 mice. The glomerulus-level analysis of rise time (Fig. 8) includes data from 10 glomeruli in 6 mice. The comparisons between tdTomato^+^ and tdTomato^−^ neurons (Figs. 6 and 8B) were carried out on fewer cells and glomeruli, those that satisfied additional criteria (see Results).

**Figure 8.**
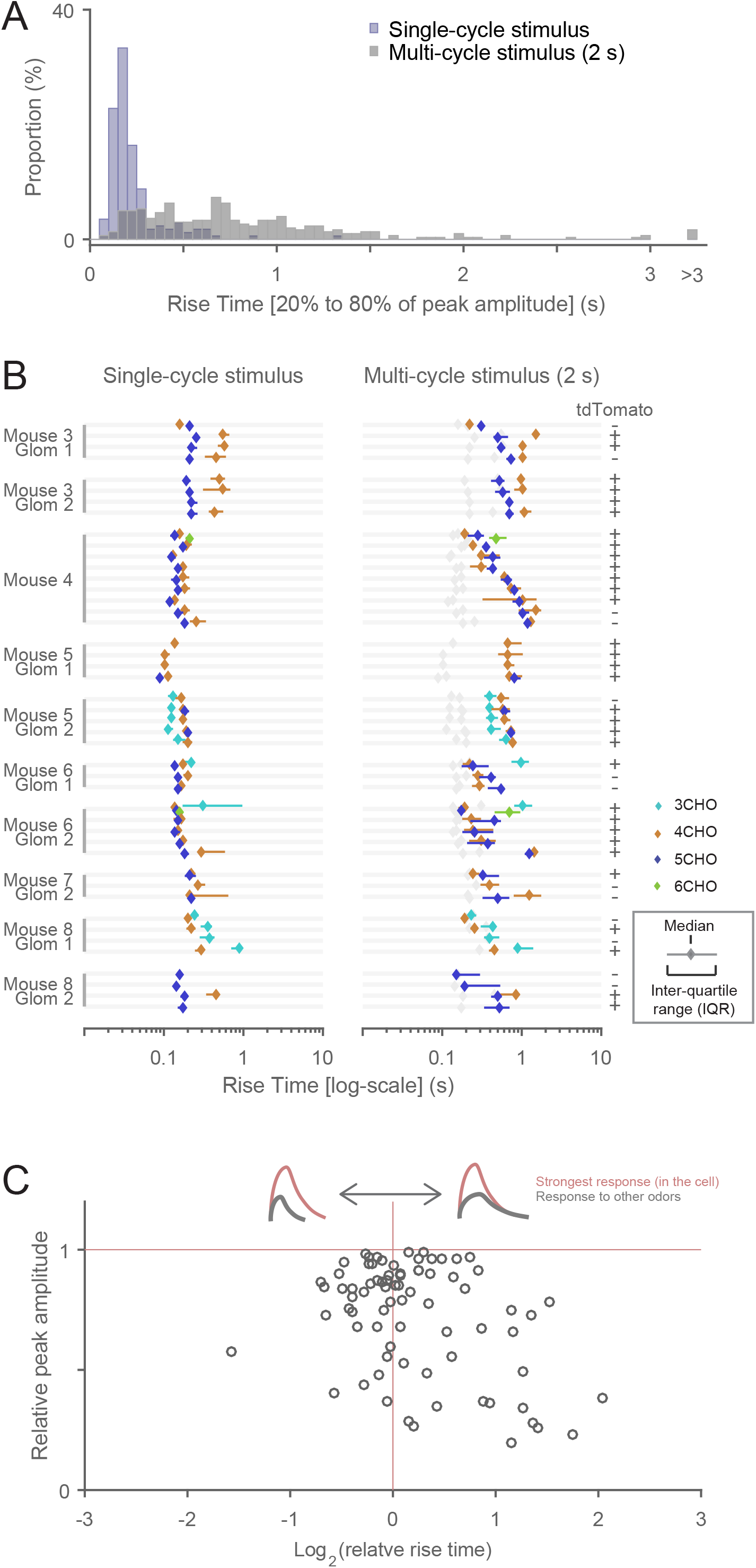
Analyses of the rise time of odor-evoked calcium transients. A. Distribution of rise times, defined as the duration between the points when a signal reached 20% and 80% of peak amplitude. Two overlapping histograms are presented. The blue histogram represents the distribution of single-cycle stimulation, and the gray one represents the distribution of multi-cycle (2 s) stimulation. B. Rise times are presented for cells putatively associated with the same glomerulus. All cases in which rise time was successfully determined in three or more cells are presented. The median is presented as a diamond, and the inter-quartile range as a horizontal bar. No horizontal bar means that the inter-quartile range is smaller than the size of the diamond marker. In the right panel (multi-cycle stimulation), the data from the left panel (single-cycle stimulation) are replicated in gray to facilitate comparison. Note the logarithmic scale on the x-axis. Expression of the GAD2 marker tdTomato (positive (+) or negative (-)) is indicated at the right. The indexes for mouse and glomerulus are shown at the left: these indexes correspond to those in Fig. 6. C. The relationship between onset latency and peak response amplitude is plotted. Because peak amplitudes can be compared only within the same cell, only data from cells in which the rise time was successfully determined for more than one odorant were used in the analysis (see text for details). Note the logarithmic scale on the x-axis.

Normality of the data is not assumed in the statistical tests. P-values less than 0.05 were considered statistically significant. The following is a list of the statistical tests used. Fig. 4D: Mann-Whitney U-test. Fig. 5B: Kruskal-Wallis test, for comparisons among glomerulus–odor pairs. Fig. 6: comparison with shuffled data (see below) for differences across glomerulus–odor pairs; Mann-Whitney U-test for the comparison of glomerulus-controlled relative onset latency (see the previous section) between tdTomato^+^ and tdTomato^−^ neurons. Fig. 7C: Wilcoxon signed-rank test. Fig. 8B: comparison with the shuffled data for differences across glomerulus–odor pairs. Mann-Whitney U-test for the comparison of glomerulus-controlled relative rise time between tdTomato^+^ and tdTomato^−^ neurons.

To statistically test whether the onset latency or rise time depended on the glomerulus–odor pairs (termed “groups” here) (Figs. 6 and 8B), the inter-quartile ranges (IQRs) of each group were compared with the corresponding values from shuffled data sets, according to the following procedure. First, the IQRs of onset latency or rise time were estimated for each group. The median IQR across all groups was chosen as the parameter to be tested. To shuffle the data, the association between the cell–odor pair and the group was randomly permuted, with maintaining the sizes of the groups (i.e., the number of neurons in each group). Shuffling was conducted for 100 000 runs, and the corresponding test parameter (median IQR of onset latency or rise time across groups) was calculated for each run. The P-value was defined as the percentage of shuffled runs in which the test parameter was smaller than that from the actual data.

## Results

To analyze temporal coordination among JG cells in the glomerular network, we primarily focused on PG cells. PG cells are monoglomerular and are thus thought to receive little or no direct input from other glomeruli (Wachowiak and Shipley, 2006; Kiyokage et al., 2010). In addition, they are the most abundant cell type in the GL and can be genetically labeled. These traits make PG cells an excellent target for studying the coordination of odor-evoked activity in the glomeruli. Rather than selectively expressing the genetically encoded calcium indicator (GECI) GCaMP6f in PG cells, we genetically labeled PG cells with a red-fluorescent protein tdTomato and expressed GCaMP6f via an injected AAV vector under the pan-neuronal synapsin promoter. This labeling strategy allowed us to record from genetically identified PG cells as well as neighboring neurons of other types.

### Expression pattern of fluorescent proteins

We used GAD2 (GAD65) as a molecular marker for a subset of PG cells (Kiyokage et al., 2010) in the GL, and genetically labeled this subset with a cre-dependent tdTomato reporter mouse (Madisen et al., 2010) that was crossed with a Gad2-IRES-Cre mouse (Taniguchi et al., 2011). First, we examined the expression pattern of tdTomato (Fig. 1). tdTomato expression was observed in the GL and all inner layers of the OB. We occasionally observed off-target tdTomato expression in a small subset of OSN axon bundles as well (data not shown). In the GL, while many JG-cell somata were labeled with tdTomato, fluorescence in the glomeruli was less prominent. Nearly all tdTomato^+^ somata were also positive for GABA immunolabeling (Fig. 1A). A previous study reported that ~37% of JG cells were positive for GAD65, ~32% were positive for GAD67, and ~16% were positive for both GAD67 and GAD65 (Parrish-Aungst et al., 2007). Therefore, we expected to observe a significant proportion of GABA^+^/tdTomato^−^ cells, corresponding to GAD67^+^/GAD65^−^ cells, but this was not the case. We speculate that some of the GAD67^+^/GAD65^−^ cells may have expressed Cre-recombinase (under the Gad2 promoter) earlier in the animal’s life and thus expressed tdTomato. In this scenario, tdTomato^+^ neurons would represent both GAD65^+^ JG cells and a subset of GAD67^+^/GAD65^−^ cells. This raises the possibility that sSA cells, a separate class of JG cells positive for GAD67 (Kiyokage et al., 2010), were included in the tdTomato^+^ cells. Therefore, we next examined the co-expression of tdTomato and tyrosine hydroxylase (TH), which is a marker of sSA cells in the OB (Kiyokage et al., 2010; Banerjee et al., 2015) (Fig. 1B). We found that TH^+^ cells included both tdTomato^+^ and tdTomato^−^ subpopulations (see white and yellow arrowheads in Fig. 1B). This result is qualitatively consistent with the report that a subpopulation of TH^+^ JG cells expresses GAD65 (Parrish-Aungst et al., 2007). In a previous study (Wachowiak et al., 2013), TH expression in JG cells was compared with GCaMP3 expression driven by the same mechanism as tdTomato in this study (a cre-dependent GCaMP3 reporter mouse crossed with the same Gad2-IRES-Cre driver mouse). They reported that 14% of TH+ cells co-expressed GCaMP3, whereas the proportion of TH+/tdTomato^+^ cells among TH^+^ cells in this study appears higher, for unknown reasons. To summarize, tdTomato expression was specific to GABAergic neurons but may not be restricted to GAD65^+^ neurons; thus we cannot rule out the possibility that a small proportion of tdTomato^+^ cells might represent sSA cells. Because we did not detect any specific difference between tdTomato^+^ and tdTomato^−^ cells in the following analyses, we did not further characterize the identity of tdTomato^+^ cells.

### Odor-evoked calcium transients in GCaMP-labeled glomeruli

At the beginning of the recording session for each mouse, we checked the positions of glomeruli that were responsive to our stimulus set by recording the odor-evoked response in a large portion of the dorsal bulb with a high-speed CCD camera (Fig. 2A). Each panel in Fig. 2A2–4 is a combination of three activity maps that represent three different time windows (see Materials and Methods for detail). The early response (0.5–2.5 s after stimulus onset) is presented in red, the responses in subsequent time windows (2.5–4.5 and 4.5–6.5 s) are presented in green and blue, respectively, and the three maps are color-merged to produce the final image. Therefore, reddish colors imply dominance of the early response and whitish colors imply long-lasting responses, for example. Consistent with previous reports (Mori et al., 2006), strong activity was elicited by aliphatic aldehydes with different carbon chains. The central region of the dorsal OB in Fig. 2A shows many globular and spatially distinct centers of activity, which each primarily represent the activity of an individual glomerulus. These glomeruli show relatively prolonged activity even after the odor stimulation (white or yellow regions). By contrast, the lateral side of the bulb (top of the images) shows activity that declines quickly after stimulus offset (red regions). There are also glomeruli that appear bluish in color (see Fig. 2A3), which represents a slow-rising calcium signal that peaked long after the stimulus offset. It is apparent that the time courses of odor-evoked calcium transients are diverse and depend on the particular glomerulus–odor pair. Of note, multiple cell types contributed to these signals from wide-field imaging because GCaMP6f was expressed under the control of the synapsin promoter. Therefore, the signals reflect not only the glomerular activity but also the activity of somata and other neuronal processes, including the inter-glomerular processes of sSA cells (Kiyokage et al., 2010) and processes in the inner layers.

### Odor-evoked calcium transients in GCaMP-labeled JG cells

For two-photon recordings of individual JG cells, we chose the recording location guided by the data from wide-field imaging. Briefly, we first identified a candidate location where we found a cluster of several glomeruli, each of which responded to our stimulus set with a discriminable odor selectivity. At the location, we first conducted conventional XYT scans to assess the odor-evoked response of glomeruli and JG cells (Fig. 2B). On the basis of the shape and “color” of the activity maps, we could reliably match the glomeruli in the two-photon imaging with those in the wide-field imaging (Fig. 2). We kept the recording depth relatively shallow (45–98 μm, 62.8 ± 16.6 μm mean ± sd, from the dura mater) so that the glomeruli were included in the field of view. Since we needed to determine the recording sites (regions of interest; ROIs) for random-access scanning prior to the recording, we next picked 15 cells on the basis of the two-photon response maps (e.g., Fig. 2B) and the unprocessed two-photon microscope images (see also the Materials and Methods). Among 120 cells from 8 recordings, 79 were identified as tdTomato^+^ (9.9 ± 2.6 per recording, mean ± sd). The tdTomato^−^ recorded cells may include GAD65^−^ PG cells, sSA cells, and ET cells. ET cells, however, may be under-represented, considering that the AAV vector we used (AAV2/1) preferentially infects PG and/or sSA cells (Adam et al., 2014) and that the somata of ET cells are located in the deeper part of the GL (Pinching and Powell, 1971; Macrides and Schneider, 1982; Hayar et al., 2004).

Examples of odor-evoked calcium transients are presented in Fig. 3A. Because the random-access recording method records data only from a set of pre-determined ROIs (see Figs. 3B and 4B), the obtained data is not a series of XY images but multiple traces from the ROIs (optical multi-ROI recording). Conventionally, odor-evoked responses are measured by presenting odors for several seconds (2 s in this study) as shown in Fig. 3A (left panel). The responses to a 2 s stimulus exhibited some diversity in their time courses. Whereas previous electrophysiological experiments demonstrated that odor-evoked responses of PG cells tend to occur as bursts of action potentials that are locked to the respiration cycle (Tan et al., 2010; Homma et al., 2013; Livneh et al., 2014), we observed respiration-locked modulation of calcium signals in only a minor subset of traces (data not shown). The typical time course of the odor-evoked calcium transients with 2 s stimulation was a monotonic rise followed by a monotonic decline, at various rates for each (Fig. 3A). However, the number of respiration cycles for the transients to reach the peak was heterogeneous. In some cases, the signal reached its peak amplitude during the first odor inhalation, while in other cases it took two or more inhalations to reach the peak. Although such heterogeneity in the response time course of JG cells is consistent with that observed in previous calcium-imaging studies (Wachowiak et al., 2013; Adam et al., 2014), this heterogeneity complicates the physiological implications of information extracted from the calcium signal (e.g., the peak amplitude). To obtain simpler and more comprehensible time courses, we used briefer stimuli that were controlled to the duration of a single respiration cycle (see Materials and Methods). The right panel of Fig. 3A shows examples of responses to such single-cycle stimuli. As expected, we observed more-stereotyped time courses with these stimuli than with multi-cycle (2 s) stimuli.

### Response-based estimation of associated glomerulus

JG cells that are associated with the same glomerulus show highly correlated stimulus selectivity (Tan et al., 2010; Kikuta et al., 2013), and are thought to be located in the proximity of the associated glomerulus (Homma et al., 2013; Kikuta et al., 2013; Adam et al., 2014). We thus wondered if we could infer the association between the recorded cells and a particular glomerulus from their odor response and the location of the cell body. By comparing the odor-evoked responses to different odors, we observed that some of the recorded cells indeed showed highly correlated response profiles (Fig. 3C). To systematically compare the odor responses, we used the data from single-cycle stimulation trials and calculated the response amplitude, defined as the integral of each ΔF/F_0_ trace (i.e., the area under the trace). The five-element response vectors (middle panel of Fig. 3C) that represent the response amplitudes to the five odors were often sufficient for comparison. However, we also considered the ten-element difference vectors that were composed of the differences between every pair of response amplitudes (right panel of Fig. 3C); these were useful to distinguish between the subsets of cells with relatively similar response vectors. Combining these data with the location of individual cells (Fig. 3B), we grouped together the cells that surrounded the same glomerulus and that showed highly correlated response profiles, and assumed that these neurons were associated with the same glomerulus (Fig. 3B2, C; see Materials and Methods for details). Note that only time-averaged data (area under the trace) were used for this glomerulus-assignment process. We deliberately avoided employing the temporal patterns of calcium transients during this process because we will discuss the dynamics of calcium transients with reference to these groups in the subsequent analyses. In the representative case shown in Fig. 3, we identified two groups of four cells each (shown in red and blue). We also found another group with two cells (shown in brown), although this group was not considered in the subsequent analyses because it contained too few cells. We were able to find 14 groups of three or more cells that satisfied the criteria in the eight fields of view from eight mice.

### Baseline activity of the GCaMP-labeled JG cells

Prior to a detailed analysis of the odor-evoked response, we briefly characterized the calcium signals in the pre-stimulus period and examined if they could potentially affect the analyses of the odor-evoked responses (Fig. 4). Fig. 4A represents examples of traces during the pre-stimulus period. Whereas previous electrophysiological recordings reported that some JG cells exhibit action potentials that are coupled to the respiration cycle even without an odor stimulus (Homma et al., 2013, Livneh et al., 2014; Najac et al., 2015), we did not observe calcium signals that were regularly coupled to the respiration cycle in our cellular recordings despite a sufficiently high sampling rate. Possible explanations may include relatively few action potentials per cycle, relatively slow kinetics of either the GECI signal or intracellular calcium itself in the cell body, which may mask the respiration-locked modulation, or a combination of all of these. In contrast to the lack of respiration cycle-locked transients, we observed occasional fluctuations of the GCaMP signal that were absent or much less prominent in the simultaneously recorded red fluorescence from (calcium-insensitive) tdTomato. To compare the size of pre-stimulus fluctuations with the size of the odor-evoked response, we calculated the ratio of the peak amplitude of the odor-evoked response to the range (the difference between the 1st and 99th percentiles) of the pre-stimulus period signal (Fig. 4C). In the vast majority of cell–odor pairs (s/n > 1 in 99.7% of the cell–odor pairs with odor-evoked responses), the odor-evoked response was much larger than the pre-stimulus fluctuation, suggesting that the pre-stimulus activity would not significantly affect the analysis of odor-evoked calcium transients.

We also noticed that signal fluctuations during the pre-stimulus period tend to be synchronized among the cells putatively associated with the same glomerulus (see previous section). This observation suggests that pre-stimulus activity is driven by some form of glomerular input or spontaneous local network activity within a glomerulus. To examine this possibility, we calculated the pairwise Pearson’s correlation coefficient between the time courses during the pre-stimulus period for all simultaneously recorded pairs of cells. The distribution of correlation coefficients is presented separately for the cells putatively from the same glomerulus and all other pairs (Fig. 4D). The correlation coefficients were significantly higher in the “same glomerulus” pairs than in the “all others” pairs (P = 2.4 × 10^−25^, Mann-Whitney U-test for the same distribution, N = 135 and 705 pairs of cell–odor pairs for the “same glomerulus” and “all others” groups, respectively). This result suggests some degree of synchrony in the pre-stimulus activity among the JG cells associated with the same glomerulus. There is a chance that the “all others” pairs may include some in which both cells were actually associated with the same glomerulus but that were mislabeled owing to our relatively conservative procedure for cell grouping.

### Glomerulus-specific onset latencies of odor-evoked calcium transients

Although it is difficult to infer the precise time course of bursts of action potentials from an odor-evoked calcium transient, it is more feasible to determine the onset latency of the transient, which reflects the onset of the burst. Therefore, we analyzed the onset latency of odor-evoked calcium (Figs. 5–7). First, we carefully determined the onset latency from our high-sampling-rate data (Fig. 5A; see Materials and Methods for details). Briefly, we approximate the earliest part of the rising signal as a linear line; the time of onset is defined as the time when this line intersects the baseline. The onset latency was defined as the difference between the time of onset and the onset of inhalation estimated from the respiration signal. Fig. 5B presents onset latencies from cell–odor pairs in which onset latencies were successfully determined in at least five trials. Although the trial-by-trial deviation was relatively large, as shown by the IQR in Fig. 5B, the median onset latency was not the same across cell–odor pairs (P = 1.8 × 10^−144^, N = 165 cell–odor pairs: Kruskal-Wallis test for non-uniform distribution). The distribution of onset latencies ranged from 0 to 150 ms. Although we do not know precisely why the shortest onset latencies were so close to the onset of inhalation (0 ms), our speculation is that the respiration signals recorded from chest movements may lag behind the inhalation at the nose. If this is indeed the case, then every onset latency reported in this study is underestimated by a certain amount.

The causes of heterogeneity in onset latency may include (1) heterogeneous inputs to different glomeruli, (2) differential interactions with the glomerular circuits, or (3) differences in cell type. Since the majority of our recorded cells were PG cells, differences in cell type are unlikely to explain all of the heterogeneity. If heterogeneous inputs to different glomeruli are the primary cause, the deviation in onset latency should be much smaller among the JG cells associated with the same glomerulus. Therefore, we examined onset latencies in the groups of cell–odor pairs putatively associated with the same glomerulus to compare the onset latencies within and across glomeruli (Fig. 6). The median onset latencies of cell–odor pairs putatively associated with the same glomerulus were distributed in relatively narrow time windows compared with the distribution as a whole, suggesting that onset latency is primarily dependent on the glomerulus (P = 2.2 × 10^−4^, comparison of in-group IQR with shuffled data). There were cases in which different odors resulted in discriminable onset latencies for cells within the same glomerulus (e.g., mouse 2 and mouse 4 in Fig. 6), implying that the onset latency depends on both the glomerulus and the odor presented. This result also rules out the possibility that the differences in onset latency are an artifact caused by the measurement error in inhalation onset.

Given that the glomerulus–odor pair was the primary determinant of onset latency, we compared onset latencies between tdTomato^+^ and tdTomato^−^ cells to examine the possible contribution of cell type to the onset latency. When we examined the relative order of onset latency within a glomerulus–odor pair (see Materials and Methods), we did not see any differences with regard to tdTomato expression (P = 0.39, N = 41 tdTomato^+^ and 26 tdTomato^−^ cell–odor pairs: Mann-Whitney U-test).

### Onset latencies of JG cells with the same odor-response profile

To gain insight into the possible role of glomerular circuits in onset latency, we next compared the onset latency among cells within the same glomerulus–odor pair (Fig. 7). The first observation that drew our attention was that the trial-by-trial deviation of each cell (summarized as the IQR) was larger than the deviation of median onset latencies across cells (Figs. 6 and 7A1–2). To understand the origin of this deviation, we re-organized the data and sorted them by trial (Fig. 7A3–4). We found that the deviation of medians across trials was much larger than the deviation of medians across cells, suggesting that the large intracell deviations are caused by the deviation across trials, which might be related, for example, to a fluctuating state in the animal or to measurement error in the inhalation onset. Of note, we did not find a clear relationship between the median onset latency within the trial and the order of acquisition (data not shown). As presented in Fig. 7C, the IQR across trials was larger than the IQR across cells for most of the glomerulus–odor pairs (P = 2.2 × 10^−4^, N = 20: Wilcoxon’s signed-rank test). Therefore, when the onset latencies were aligned to the median of the corresponding trial, the deviation of each cell across trials substantially decreased (Fig. 7B). In the case presented in Fig. 7B, the order of the aligned onset latencies was well preserved between the two stimuli (note that the same color represents the same cell). Although this may imply the presence of a systematic difference in onset latency among these cells, examining this further would require a larger data set than we currently have, with multiple stimuli and more cells.

### Distribution of the rise time in single-cycle stimulation

In addition to onset latency, an odor-evoked burst of action potentials can be characterized by, for example, the duration and the number of spikes. With the assumption that the duration of the rising phase of a calcium transient (rise time) is closely correlated with the duration of burst activity, we analyzed the rise time of odor-evoked calcium transients (Fig. 8). We defined the rise time as the duration for a calcium signal to rise from 20% to 80% of the peak amplitude. Because we could not distinguish individual respiration cycles in the calcium transients evoked by multi-cycle stimulation, we focused primarily on data from single-cycle stimulation.

First, we simply looked at the distribution of rise times across odor–cell pairs (Fig. 8A). As expected, the large majority (84.8%) of rise times for single-cycle stimulation (blue) were shorter than 300 ms, suggesting that the action potentials halted (or greatly decreased) within a single respiration cycle. (Typical breathing rates during the recordings were 120–200 cycles/min, or 300–500 ms/cycle.) The presence of relatively short rise times may indicate spike activity of shorter duration in these cells. A minor proportion had longer rise times, likely reflecting the persistence of activity even after the stimulus offset. By contrast, the rise times under 2 s stimulation (gray) were distributed more broadly, reflecting the heterogeneous time courses of calcium transients in this condition (Fig. 3A). Next, as for onset latency (Fig. 6), we grouped the data from cell–odor pairs putatively associated with the same glomerulus, to examine the contribution of glomerular inputs (Fig. 8B). With single-cycle stimulation, rise times depended on the glomerulus–odor pair with statistical significance (P < 10^−5^, comparison with shuffled data, 62 cell–odor pairs from 12 glomerulus–odor pairs with 4 or more neurons). This was not the case for the multi-cycle stimulation (P = 0.12). In Fig. 8B, the IQRs for each cell–odor pair (horizontal bars) were not very different from the deviation among median rise times within the same group. This was in clear contrast with the results for onset latency (Figs. 6 and 7A) and suggests little trial-by-trial deviation (see above).

As for onset latency, above, we compared the relative rise times of tdTomato^+^ and tdTomato^−^ neurons putatively associated with the same glomerulus and did not see any statistically significant differences (P = 0.28 for single-cycle stimulation, P = 0.39 for multi-cycle stimulation, N = 30 tdTomato^+^ and 14 tdTomato^−^ cell–odor pairs from eight glomerulus–odor pairs that included four or more neurons: Mann-Whitney U-test). Finally, we examined the correlation between transient rise times and peak amplitudes, as a strong correlation between these would not be compatible with our assumption that rise time reflects the duration of burst activity. We only considered data from single-cycle stimulation in this analysis. Since we cannot compare response amplitudes across different cells, we analyzed cells that were activated by more than one odorant. For each cell, all rise times and response amplitudes were normalized to those of the response with the largest peak amplitude (the reference response), and plotted in Fig. 8C (reference responses are omitted but would appear at [0, 1] on the plot). Among 77 cell–odor pairs, the onset latency was shorter in 34 cell–odor pairs and longer in 43 cell–odor pairs than the corresponding reference response. Thus, it is highly unlikely that rise time is accounted for solely by the response amplitude, which favors our assumption that rise time is correlated with the duration of burst activity.

## Discussion

In this study, we recorded odor-evoked calcium transients from individual JG cells in anesthetized free-breathing mice at an extremely high sampling rate and measured the onset latency and rise time of the transients. We examined these properties in sets of JG cells putatively associated with the same glomerulus (“homoglomerular” JG cells). While the onset latency of JG cells spans ~150 ms when glomerular association is not taken into account, the range of onset latencies is substantially smaller (typically a few tens of milliseconds) among homoglomerular JG cells. Similar glomerulus specificity was also found for rise times when we presented a brief odor stimulus. These observations suggest that the glomeruli (more strictly, the glomerulus–odor pairs) determine a significant portion of the response time course in JG cells.

### High-speed calcium recording by random-access scanning

Here, we briefly discuss the technical aspects of this study. The intracellular calcium level has become a popular proxy for neuronal activity (Grienberger and Konnerth, 2012). It is well acknowledged that the calcium signal is much slower than the related action potentials, but this has not been a serious issue in two-photon imaging studies partly owing to the low frame rates of conventional two-photon microscopes. At the same time, such a situation has limited the analyses possible from readouts of the temporal patterns of calcium signals *in vivo*. In this study, we used a random-access scanning technique (Iyer et al., 2006; Grewe et al., 2010) and sampled the calcium transients at a high rate (667 Hz). This high sampling rate was achieved by limiting the number of pixels sampled. Thus, the sampling duration *per pixel*, which often limits the s/n ratio of two-photon recording, was no different from the full-frame scan. While random-access scanning has tended to be utilized more for volumetric recordings of numerous cells (Göbel et al., 2007; Froudarakis et al., 2014), high-fidelity recording is another powerful application of the technique (Grewe et al., 2010). Technically, our scanner is capable of scanning at *per-pixel* sampling rates up to ~10 times higher than in this study (with lower s/n ratio). Thus, this technique is compatible with two-photon voltage recording of action potentials with voltage-sensitive fluorescence proteins, which have been continuously improving in efficacy (Lin and Schnitzer, 2016; Chamberland et al., 2017).

### Heterogeneity in the dynamics of odor-evoked calcium transients

In experiments with anesthetized free-breathing animals, it is common practice to present an odor stimulus for several seconds, lasting over multiple respiration cycles. In our recording of odor-evoked calcium transients in the GCaMP6f-expressing JG cells (dominated by PG cells), it took a different number of respiration cycles to reach the transient peak in different cells (or cell–odor pairs); this is also illustrated by the heterogeneity in rise times (the period from 20% to 80% of the peak signal amplitude; Fig. 8A, B). It is not clear to what extent this heterogeneity is due to the spike–calcium relationship (Grienberger and Konnerth, 2012) or to the temporal pattern of the spikes. The phenomenon may potentially be understood better by inferring the action potentials from the calcium signals (Yaksi and Friedrich, 2006; Theis et al., 2016; Rahmati et al., 2018). In this study, we did not take this path because of our lack of easy means to validate the inference. It is indeed an interesting question whether such methods could successfully reconstruct the action potentials in this case: reconstruction may be challenging in PG cells owing to the respiration-coupled bursting activity that may make the spike–calcium relationship strongly non-linear.

Such heterogeneous time courses also raise a practical problem for quantifying the amplitude of calcium transients. For example, peak amplitude is often used to quantify the amplitude of transients, but it may be associated with different numbers of respiration cycles across cells. Note that this issue is also relevant to the analysis of rise time. In this study, we used single-respiration-cycle stimuli to prevent signals from accumulating over multiple respiration cycles for analyses involving comparisons of the response amplitude (Figs. 3 and 8C). Although we do not have enough data to address whether the problem is also relevant to other types of neurons in the olfactory system, diverse calcium-transient time courses have been reported in multiple cell types (Wachowiak et al., 2013). Thus, there may be other cases where the shape of the calcium transients deserves consideration during the quantification of response properties.

### Glomerular circuits and glomerulus-specific onset latency

The glomerulus-specific onset latencies of JG cells are characterized by relatively small deviations in onset latency within the same glomerulus and larger deviations across glomeruli. The most plausible explanation for this is that the JG cells follow the onset of sensory inputs in each glomerulus, as the onset latency of sensory inputs from OSNs is heterogeneous across glomeruli (Spors et al., 2006; Carey et al., 2009). On the other hand, the small deviations within a glomerulus may imply that the sensory inputs from many OSNs are accumulated in such a time window, given that an axon from each OSN covers a small fraction of a glomerulus (Hálasz and Greer, 1993). It is worth noting that most of our analyses were limited to relatively strong responses as we only used data with a high s/n ratio. It is thus an interesting question whether a weaker stimulus would broaden the distribution of onset latencies among homoglomerular JG cells. If this was indeed the case, such a mechanism could play an important role in the intraglomerular circuit, where many JG cells of the same type (or subtype) receive homogeneous inputs and perform the same function (e.g., feedforward excitation/inhibition). For instance, even if a class of JG cells collectively releases the same total amount of neurotransmitter to the glomerular circuit, the consequences would be different if highly synchronized neurons release the transmitter at a high density over a short time window, versus if less-synchronized neurons release the transmitter at a lower density over a longer time window.

### Implications for the function of the OB and glomeruli

Although our recordings were limited to JG cells, the glomerulus-specific response time course of these cells suggests that the time course of sensory inputs is highly glomerulus-specific. Each glomerulus was driven at a different time, with a strong input in the first tens of milliseconds. The temporal pattern of odor-evoked activity has been shown to be a part of stimulus representation in the principal neurons (Cury and Uchida, 2010; Uchida et al., 2014). Our results support the idea that sensory inputs play a role in the temporal coding of odor stimuli (Schaefer and Margrie, 2007; Raman et al., 2010). For example, a coarse temporal pattern may be determined by the sensory inputs (as well as the glomerular circuits) and be largely homogeneous among homoglomerular mitral cells. The odor-evoked activity, including the temporal pattern, may then be refined in subsequent OB circuits so that it is diversified among individual principal neurons (Dhawale et al., 2010; Kikuta et al., 2013). Furthermore, glomerulus-specific onset latencies may play a role in the horizontal connections among glomeruli (Aungst et al., 2003; Kiyokage et al., 2010; Banerjee et al., 2015). Early responding glomeruli may be capable of modulating the activity in neighboring glomeruli more strongly, although the computational advantage of such a mechanism is not obvious in a system that supposedly lacks a topographic representation of stimulus space (Cleland and Sethupathy, 2006; Cleland, 2014).

## Acknowledgments

We thank Wei Chen for his contribution to the setup of the acousto-optic deflector two-photon microscope. S.N. is supported by NIH/NIDCD Grant R01DC013802. F.I. is supported by NIH/NIDCD Grant R01DC016307. S.Z. and X.L. are supported by NSFC Grant 81327802.

